## Supplementary material for "Transcriptome and Chromatin Landscape of iNKT cells are Shaped by Subset Differentiation and Antigen Exposure": Four supplementary figures

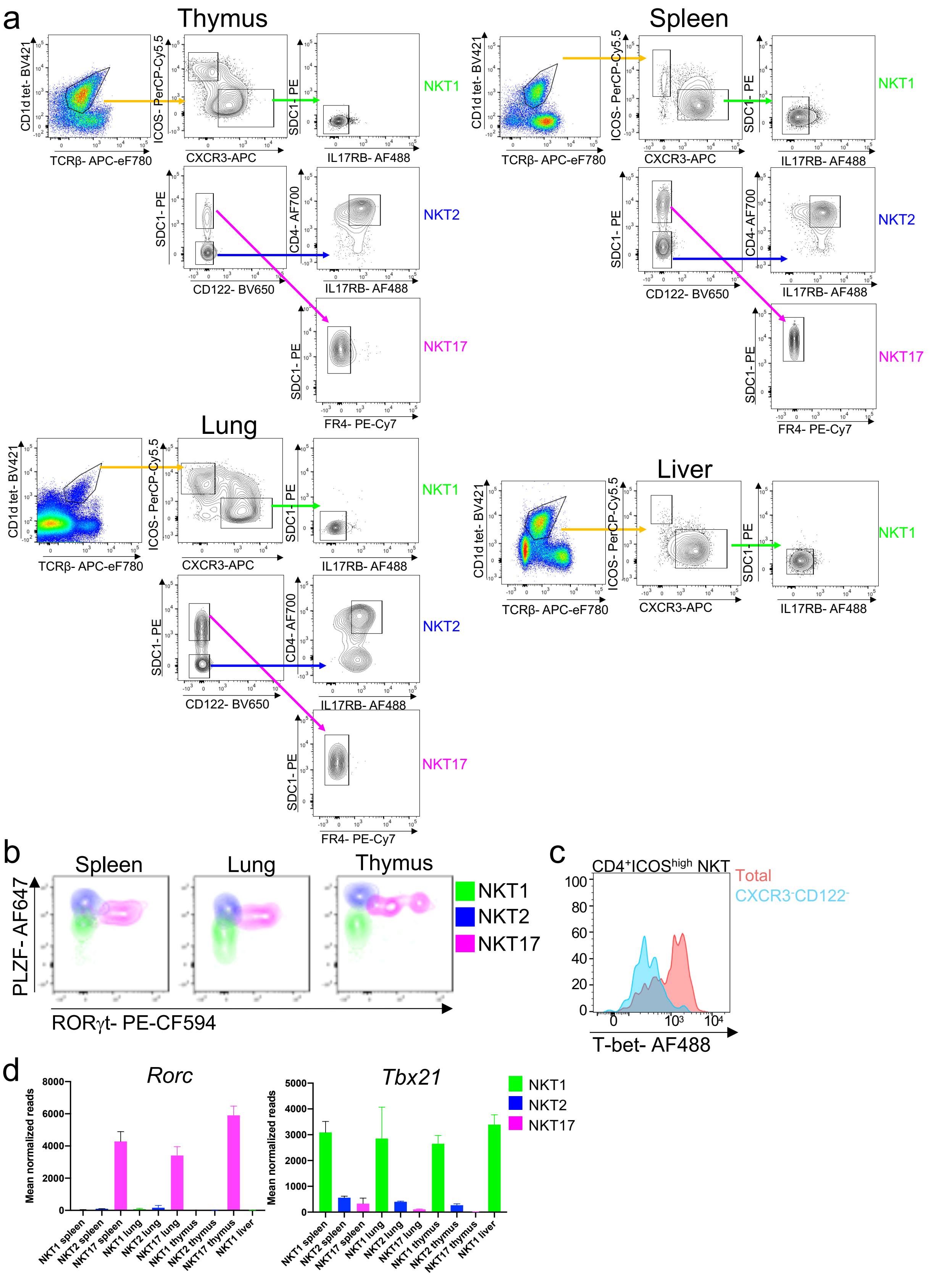


**Supplementary Fig. 1. iNKT cell subset gating.** a. Example of gating for isolating iNKT cell subsets from lymphocyte- and live-gated, cell suspensions enriched for *i*NKT cells as described in Materials and Methods. Note that some ICOS$^{high}$ cells express relatively high levels of CXCR3. Representative examples shown are from the thymus, spleen, lung and liver; one of five independent sorts. b. Expression of transcription factors PLZF and ROR$\gamma$t as measured by representative flow cytometry in spleen, lung, and thymus CD1d-$\alpha$galcer-tetramer-binding, TCR$\beta^{+}$ cells selected to be NKT1(CXCR3$^{+}$, ICOS$^{low}$, SDC1$^{-}$), NKT2 (CXCR3$^{-}$, ICOS$^{high}$, CD4$^{+}$,SDC1$^{-}$), and NKT17 (CXCR3$^{-}$, ICOS$^{high}$, CD4$^{-}$, SDC1$^{+}$). Results are representative of cell suspensions from two mice stained with this exact panel. The staining of all of the panel reagents was confirmed in samples from at least eight mice examined in at least four independent experiments. c. Histogram overlay of staining for T-bet in CD4$^{+}$, ICOS$^{high}$ iNKT splenocytes with removal (cyan) or without removal (pink) of cells staining with antibodies specific for CXCR3 or CD122, which were combined into one channel to facilitate simultaneous staining for both cell surface markers and transcription factors to define iNKT cell subsets and identify more uniformly a T-bet$^{low}$ population. Data are representative of samples from fourteen mice examined in seven independent experiments. d. Mean normalized reads for *Rorc* and *Tbx21* as determined by RNA-seq for cells sorted as described in Fig. Supp. 1a, determined from four to seven samples obtained from three to five sorts.

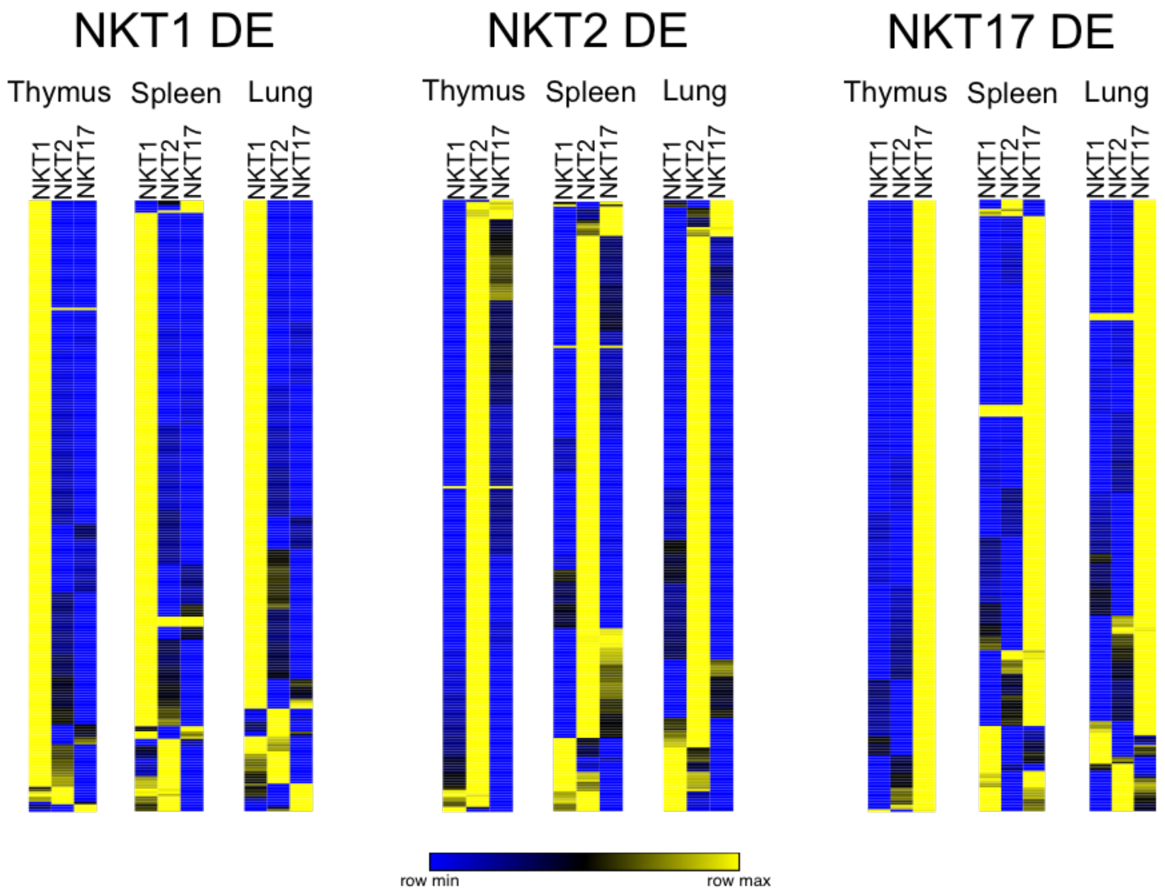


**Supplementary Fig. 2. Reproducibility of thymic iNKT cell subset gene signatures.** Thymic iNKT cell subset transcriptomic signatures, as defined in Engel *et al*. 2016, were compared to the data in this study. Differentially expressed (DE) genes, comparing one subset to the other two, n > 2-fold difference, adjusted p = 0.05, from the earlier study were compared to the results from this study.

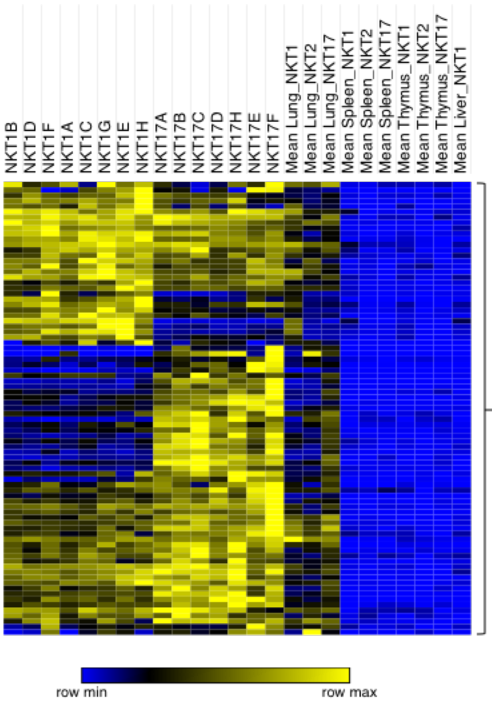


2810459M11Rik
2900060B14Rik
4930417O13Rik
5430416N02Rik
Abcb1b
Adrb2
Arid5a
Arl5b
Arl5c
Arrdc3
Atf3
Atf4
Atp2b1
Bambi
Bcl2a1b
Bcl2l11
Btg2
C920009B18Rik
Cav1
Ccrl2
Cd69
Cdkn1a
Chrne
Cldn5
Clk1
Cpne8
Csrnp1
Ctla2a
Ctla4
Cwc25
Dennd4a
Dgat1
Dnajb4
Dnajb9
Dusp5
Emp1
Ets2
Fasl
Fos
Fosb
Fosl2
G530011O06Rik
Gabpb1
Gadd45b
Gm11974
Gm11978
Gm19757
Gna13
Got1
Gpr116
Gpr132
Gprc5a
Gramd3
H3f3b
Hdc
Hey1
Hilpda
Hist1h1c
Hist1h2b
c
Hspa1a
Hspa1b
Ier5l
Ifngr2
Ifrd1
Il1rl1
Ipmk
Irg1
Isca1
Isy1
Kctd12
Kdm6b
Klf4
Klf6
Lmn
Malt1
Map1lc3a
Metrnl
Muc1
Mxd1
Naa38
Neurl3
Nfil3
Nfkbia
Nfkbid
Nfkbiz
Nr4a1
Nr4a2
Nr4a3
Obfc2a
Odc1
P2ry10
Pard6a
Pde4d
Per1
Pgm2l1
Phlda1
Pim1
Plaur
Plcxd2
Plk2
Pmp22
Polg2
Prdm2
Rab4a
Ramp3
Rgs1
Rnf125
Rnf19a
Rrad
Samd11
Samd8
Samsn1
Scgb1a1
Serinc1
Serpinb6b
Sgip1
Sgtb
Sik1
Snx18
Spty2d1
Sult2b1
Tgif1
Thbd
Tigit
Tinf2
Tiparp
Tmem64
Tnfrsf10b
Tnfrsf1b
Tob1
Tob2
Trim43b
Trmt61b
Trp53inp2
Tsc22d3
Tspan7
Tubb2a
Vgll4
Zbtb10
Zc3h12a
Zc3hav1
Zfp36l2

**Supplementary Fig. 3. iNKT cell lung signature genes.** a. Heat map of normalized read counts from RNA-seq for genes with significantly elevated reads (raw p value of <0.1) in lung samples in all pairwise comparisons between lung and other tissue samples for each iNKT cell subset. Included are the mean normalized reads of NKT1 and NKT17 cells prepared from lungs of individual mice, designated with letters A-H, and from pooled mice (4-7 experimental replicates). Alphabetical list of lung signature genes (right).

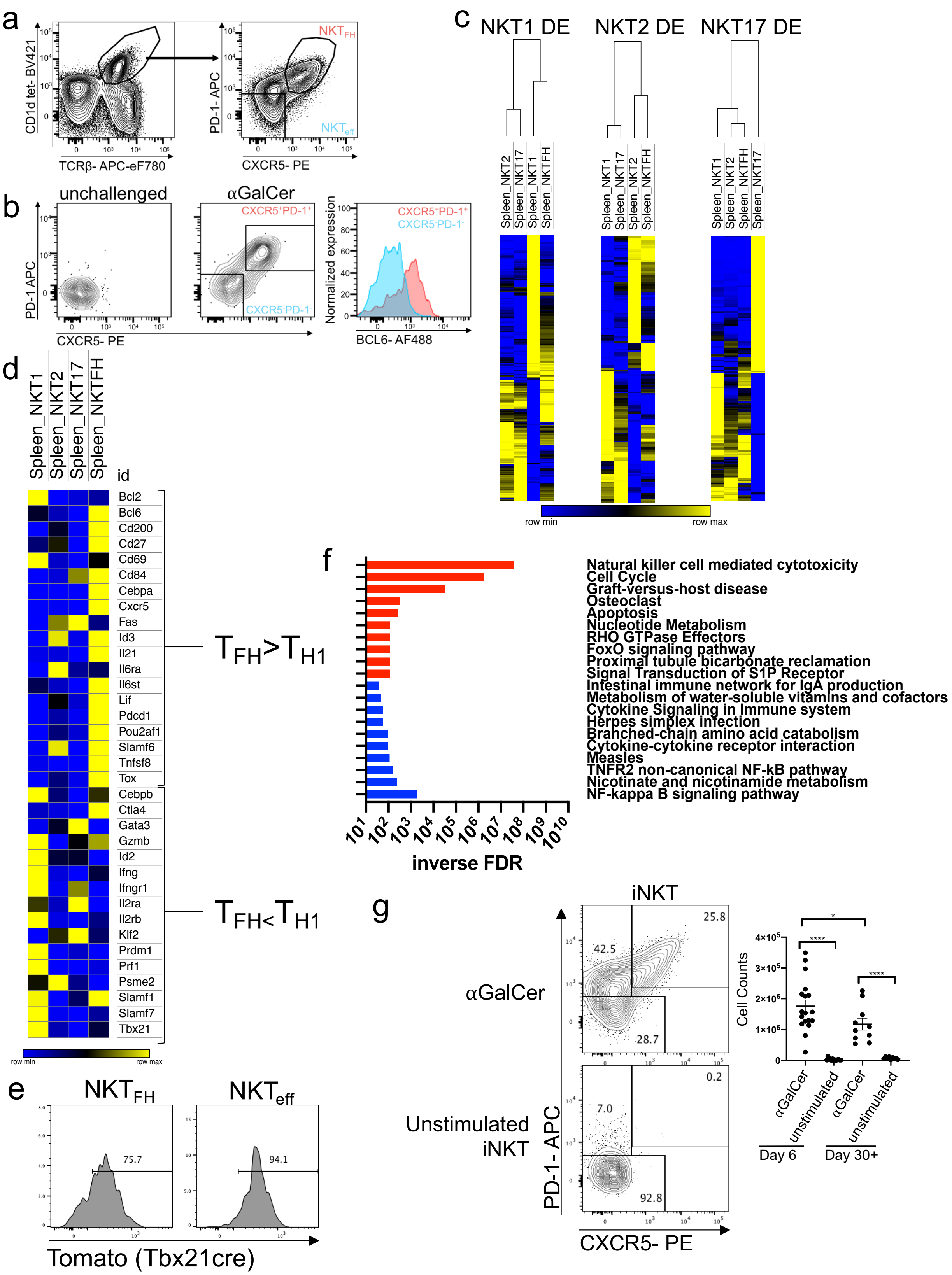


**Supplementary Fig. 4. Phenotype and transcriptome of NKT$_{FH}$ cells.** a. Splenocytes from 7-week-old C57BL/6J females injected with αGalCer 6 days prior to analysis were enriched for iNKT cells and stained as described in Materials and Methods. Plots show the staining of live, lymphocyte-gated cells with anti-TCRβ and

$\alpha$GalCer loaded CD1d tetramers (left), and the staining of CD1d-tetramer$^+$ TCR $\beta^+$ cells for CXCR5 and PD-1 (right). b. Staining for expression of PD-1 and CXCR5 in gated iNKT cells from the spleen of unchallenged mice (left panel) and $\alpha$GalCer injected mice (middle panel). Right panel: Staining for BCL6 in gated iNKT cells from immunized mice that were putative NKT$_{FH}$, or CXCR5$^+$ PD-1$^+$ (pink histogram) or that were double negative for these markers (NKT$_{eff}$, blue histogram). Data are representative of samples from 12 mice analyzed in three independent experiments. c. Heat map depicting the relative RNA-seq normalized read counts in splenic NKT1, NKT2, NKT17 and NKT$_{FH}$ cells of genes that were DE in either NKT1 (left), NKT2 (center) or NKT17 cells (right) (Padj <0.1, shrunken lfc >1 or <-1). d. Heat map of RNA-seq normalized read counts from selected genes known to be DE in T$_{FH}$ compared to T$_{H1}$ cells. e. Expression of reporter in T-bet fate-mapping mice by NKT$_{FH}$ and NKT$_{eff}$ cells 3 days-post antigen exposure. f. ConsensusPathDB gene clustering: red up, blue down. g. NKT$_{FH}$ persist in the spleen for at least one month. Left: representative flow cytometry showing NKT$_{FH}$ (TCR $\beta^+$ CD1d-tetramer$^+$CXCR5$^+$PD-1$^+$) at d30+ after $\alpha$GalCer plus unstimulated controls. Right: number of NKT$_{FH}$ at day 6 or day 30+ after $\alpha$GalCer plus controls. Data are combined from 14 experiments, n = 18 (d6 $\alpha$GalCer), n = 13 (d6 unstimulated), n = 10 (d30 $\alpha$GalCer), n = 9 (d30+ unstimulated), error bars depict SEM. Mann-Whitney test (two-tailed), $^{****}$ = p values of <0.0001, $^*$ = p value of 0.04.
